# Conservation of animal genome structure is the exception not the rule

**DOI:** 10.1101/2024.08.02.606322

**Authors:** Thomas D. Lewin, Isabel Jiah-Yih Liao, Yi-Jyun Luo

## Abstract

Species from diverse animal lineages have retained groups of orthologous genes together on the same chromosomes for over half a billion years since their last common ancestor. However, by examining interchromosomal rearrangements across all major bilaterian groups, we show that cases of high-fidelity genome structure conservation are unexpectedly rare. Large-scale genome restructuring events are pervasive, correlate with increased rates of protein sequence evolution, and may contribute to adaptation and animal diversity.

## Main text

A remarkable feature of animal genomes is the retention of orthologous genes on the same chromosomes over long evolutionary distances^1,2^. All bilaterians inherited 24 ancestral linkage groups (ALGs), sets of genes colocated on the same chromosome in their common ancestor^3^. Using orthologous genes to compare genomes, we can observe that lineages that have been diverging for over 550 million years have retained largely the same chromosomal organization, and genome structures of distantly related living species such as polychaetes, scallops, and even chordates like amphioxus differ only by a few lineage-specific chromosome fusion events^3–6^. This evolutionarily deep emergence and stability suggests that strong selective pressures have favored genome structure conservation for hundreds of millions of years.

However, this conservation of gene linkages (or ‘synteny’) can be degraded by interchromosomal rearrangements like chromosome fusion, fission, and translocation^3^. Clitellates^6,7^, nematodes^4^, planarians^8^, and bryozoans^9^ have all lost the inherited bilaterian genome structure to varying extents. Despite this, there has remained an assumption that such cases are the exception and that most bilaterian lineages retain the 24 linkage groups inherited from their common ancestor with relatively few changes, mostly involving chromosome fusions^3,7^.

In this study, we aimed to test this assumption by performing a quantitative profiling of the extent of genome structure conservation across bilaterians. We first assembled a dataset of 64 chromosome-level genomes from 15 bilaterian phyla and at least 52 classes, with the aim of maximizing phylogenetic coverage while minimizing biases caused by taxon overrepresentation. In each genome, we assigned genes to one of the 24 bilaterian ALGs using orthology and calculated a rearrangement index (RI) to quantify ALG conservation^6^. The RI quantifies ALG combining (e.g. by chromosome fusion) and splitting (e.g. by chromosome fission) and ranges from 0 (exact ALG conservation) to ≈1 (complete ALG loss). We validated this index by applying it to species with varying levels of interchromosomal rearrangements reported in the literature. RI scores were consistent with the extent of described rearrangements, confirming it as a valid metric for measuring genome structural changes (Figure S1A). We also tested the hypothesis that bilaterians can be split into two distinct groups, those with high (RI > 0.75) and low (RI < 0.75) rearrangement genomes^6^. Principal component analysis of ALG combining and splitting indices supports this division and suggests that the main factor distinguishing the two groups is increased levels of ALG splitting in high rearrangement genomes (Figure S1B–D).

Having validated the RI, we used it to quantify the level of rearrangement in each assembly in our dataset. Remarkably, of the 64 genomes analyzed, 44 are highly rearranged. Moreover, eight of the 15 phyla in our dataset contain exclusively high rearrangement genomes (Arthropoda, Bryozoa, Platyhelminthes, Nematoda, Nematomorpha, Rotifera, Tardigrada, and Xenacoelomorpha); three (Annelida, Chordata, and Mollusca) contain both high and low rearrangement genomes; and only four (Brachiopoda, Echinodermata, Hemichordata, and Nemertea) contain exclusively low rearrangement genomes (Figure 1A). The discovery that two-thirds of species in a broad sample of bilaterians have highly rearranged genomes and that over 70% of phyla contain highly rearranged species demonstrates that loss of synteny is much more common within bilaterians than previously thought.

**Figure 1.**
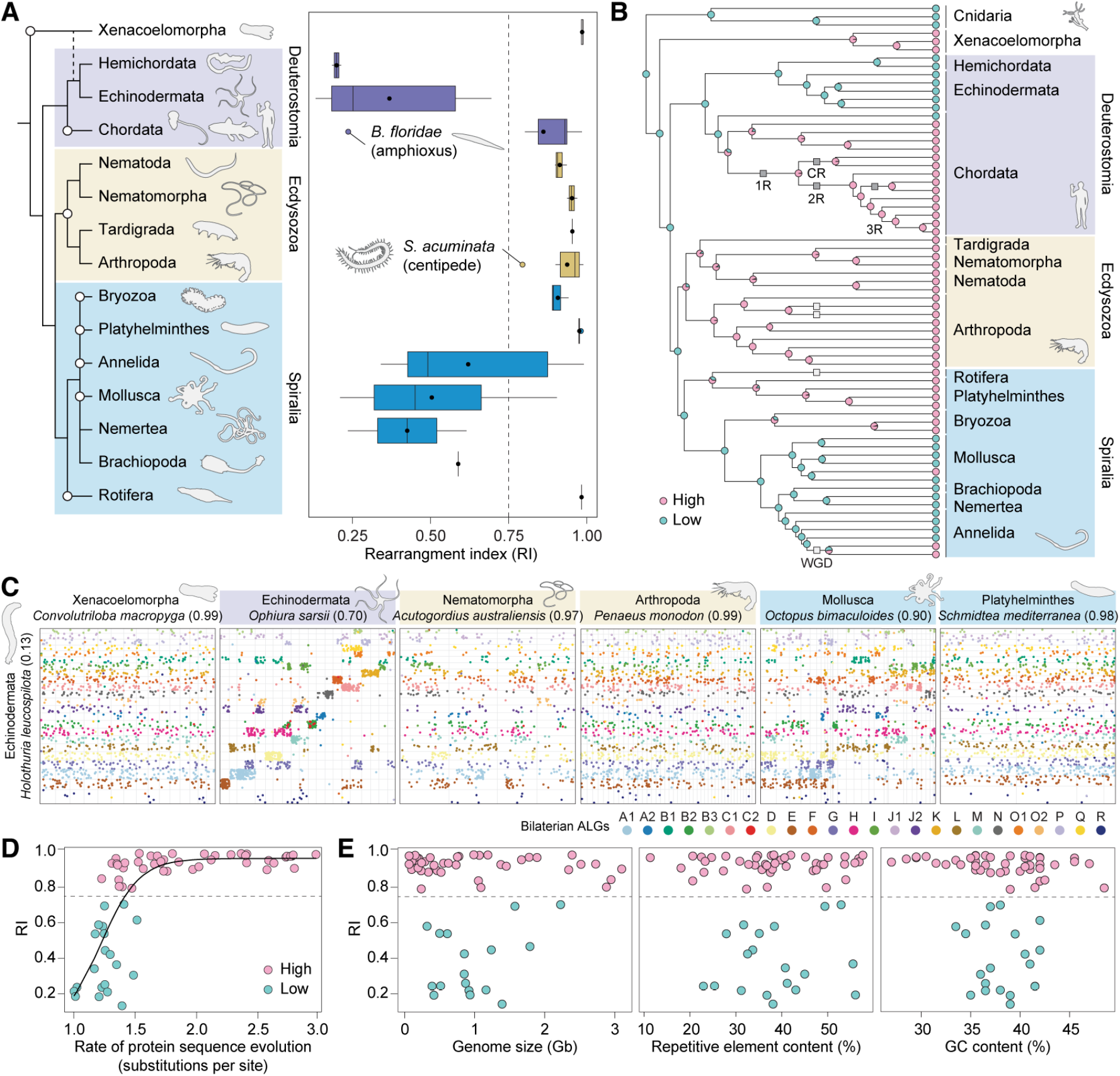
Massive genome rearrangements are the norm in bilaterians and are associated with higher rates of protein sequence evolution. (A) Boxplot of RI for 64 bilaterian species from 15 phyla. White circles indicate clades containing species with RI > 0.75. (B) Ancestral state reconstruction of genome rearrangement levels in a phylogenetic analysis of 67 animal species. Pie charts at inner nodes represent the proportion of stochastic character maps that produced each state. Maximum likelihood tree was built using 148 orthologous protein sequences. Squares indicate whole genome duplication (WGD) events. (C) Dot plots for the genome with the highest RI value (shown in parentheses) from selected phyla. Dots represent the position of orthologous genes, color-coded by their respective ALGs; chromosomes are separated by gray lines. (D) Genomes with high rates of protein sequence evolution also have high levels of interchromosomal rearrangement. Relationship was analyzed using a logistic regression model, with a residual sum of squares of 1.827. (E) There is no clear relationship between genome size, repeat content, or GC content and RI.

We next questioned whether high rearrangement genomes evolved just once or independently multiple times. Ancestral state reconstruction reveals that transitions to a high rearrangement state occurred within all three major bilaterian groups and on a minimum of seven occasions (within Chordata, Annelida, and Mollusca; in stem Xenaceolomorpha and Ecdysozoa; and at least twice in Bryozoa, Rotifera, and Platyhelminthes) (Figure 1B). All species within Ecdysozoa possess highly rearranged genomes, with ancestral state reconstruction supporting a loss of synteny common to many ecdysozoan lineages (Figure 1A,B), though chromosome-level genomes from Kinorhyncha, Priapulida, and Loricifera are needed to test this hypothesis. In some phyla, massive genome rearrangements have occurred within single classes (Annelida: Clitellata; Mollusca: Cephalopoda), while most lineages retain highly conserved genome structures (Figure S2). Overall, fourteen of 15 phyla (all except Hemichordata) have at least one species with an RI over 0.55 (Figure 1C). The dispersal of highly rearranged genomes around the bilaterian tree of life demonstrates that this state has evolved independently on many occasions in diverse lineages and at varied taxonomic levels.

Finally, we tested whether any genomic characters were predictive of RI scores. Notably, we found a strong positive correlation of RI with the rate of protein sequence evolution (Figure 1D). Lineages with whole genome duplications have typically lost the conserved bilaterian genome structure (Figure 1B), but we found no correlation between RI and genome size or repetitive element content (Figure 1E). Overall, lineages with high rates of genomic scrambling also have fast-evolving protein-coding sequences, suggesting that similar selective pressures may drive the rapid evolution of coding sequences and genome rearrangements.

Though we aimed to maximize phylogenetic coverage given current genome availability, our dataset represents a small snapshot of bilaterian diversity. Over half of the phyla remain unrepresented due to the absence of chromosome-level genomes, and Brachiopoda, Rotifera, and Tardigrada are represented by only one species. The seven transitions to a high rearrangement state identified, therefore, likely represent a significant underestimate, and this number will rise as genomes of currently unrepresented taxa are made available.

Overall, our findings reveal that extensive rearrangement of bilaterian chromosomes and subsequent erosion of synteny has occurred many times independently across the animal tree of life. In light of this, we argue that the synteny conservation reported in some phyla is the exception rather than the rule and that large-scale interchromosomal rearrangements have been pervasive throughout bilaterian evolution. This presents a startling paradox: the ancestral bilaterian genome structure has been heavily modified or completely lost in most lineages but, in a minority, has been impeccably conserved for over half a billion years. Why such different modes of genome structure are observed is, therefore, a pressing question in evolutionary genomics.

## Supplemental information

Supplemental information includes two figures, supplemental experimental procedures, data availability, author contributions, and supplemental references, and can be found with this article online.

## Acknowledgements

This work was supported by an Academia Sinica Career Development Award (AS-CDA-112-L06) and a National Science and Technology Council Research Project Grant (113-2311-B-001-026) to Y.-J.L.

