## Supplemental Information for "Conservation of animal genome structure is the exception not the rule"

### Figure S1

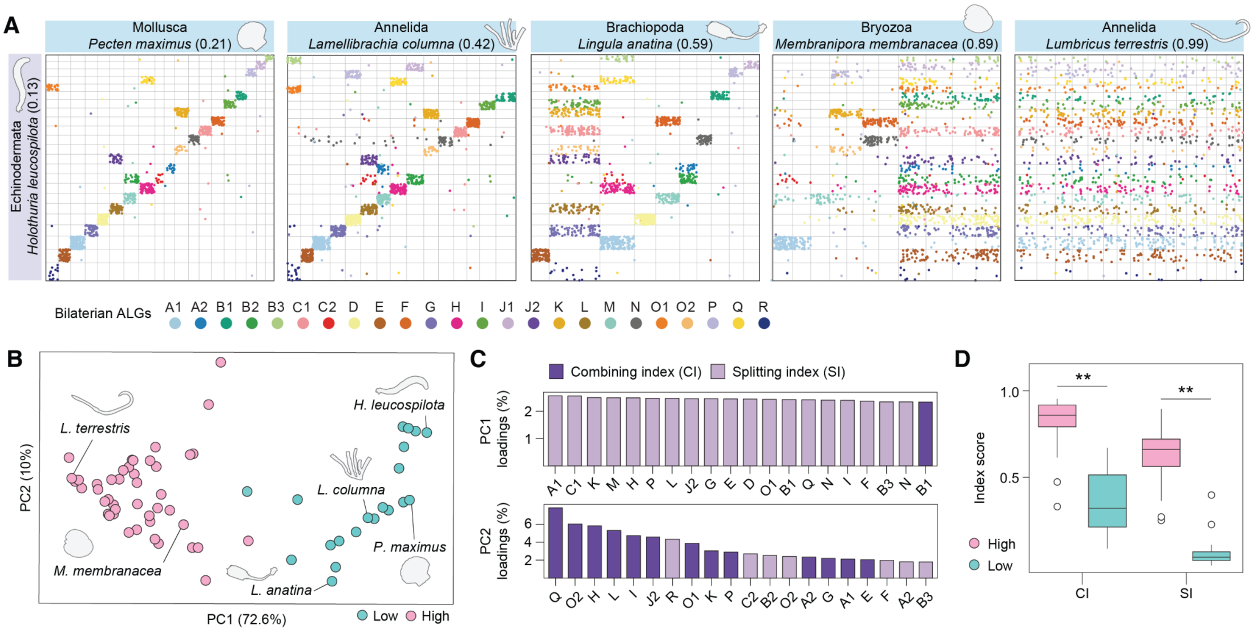


**Figure S1. Synteny quantification with a rearrangement index (RI) identifies high and low rearrangement genomes within bilaterians.**

(A) Validation of the RI[1] using genomes with known rearrangement levels. RI scores (parentheses) reflect rearrangement levels previously reported in the literature[1–5]. Each box represents a pairwise comparison of two genomes. Dots represent the position of orthologous genes in each genome, colored by their bilaterian ALG. Chromosomes are separated by gray lines. In each case, the most conserved species in the dataset (sea cucumber *Holothuria leucospilota*, RI = 0.13) is on the Y-axis. *H. leucospilota* has 23 chromosomes; 22 of which represent single bilaterian ALGs and one of which is a fusion-with-mixing event between ALGs B2 and C2. (B) PCA of combining and splitting indices for the 24 individual ALGs separates high and low rearrangement genomes. (C) Contribution of the top 20 variables to principal components shows that high rearrangement genomes have higher levels of ALG splitting. (D) High rearrangement species have significantly higher combining ($p = 2.41 \times 10^{-11}$) and splitting ($p = 2.07 \times 10^{-25}$) indices.

### Figure S2

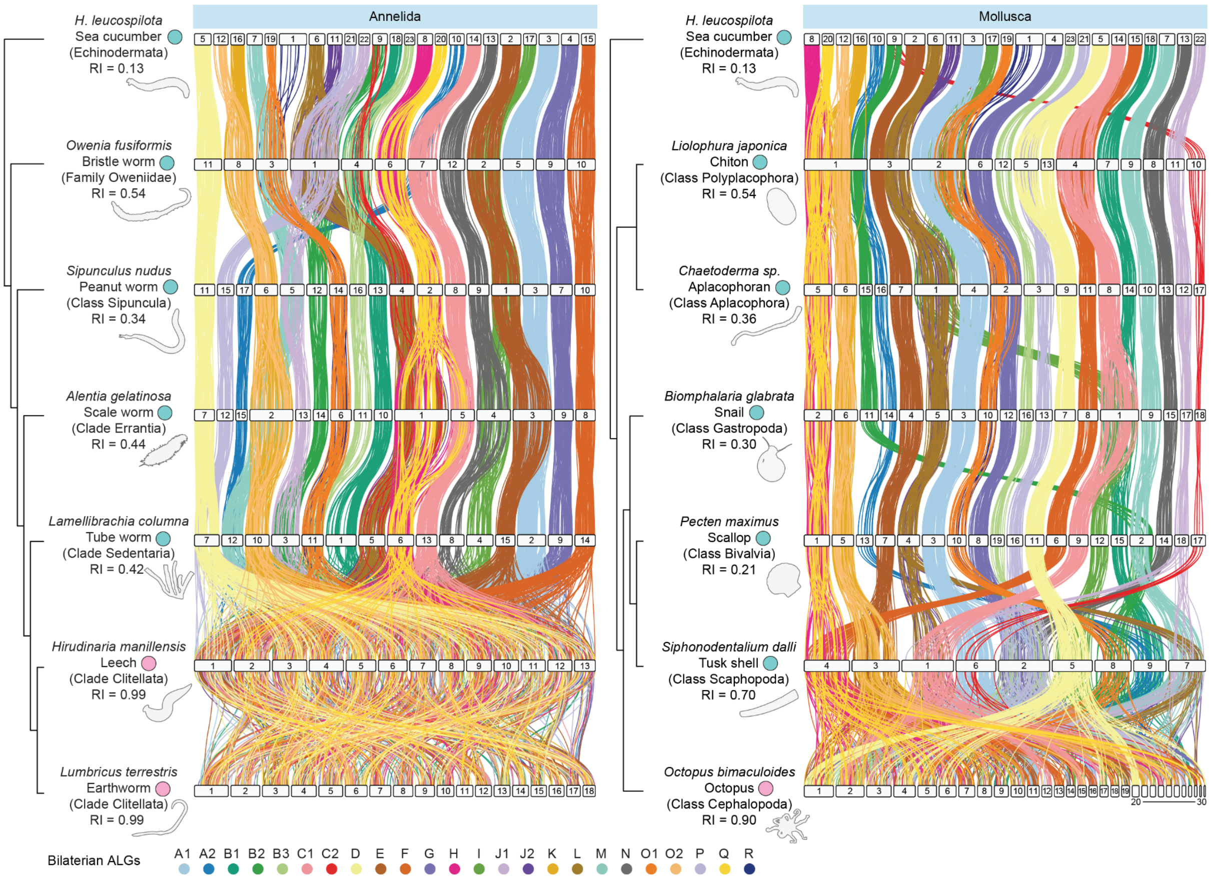


**Figure S2. Loss of bilaterian synteny can occur in specific lineages within phyla.**
Idiogram plots for annelids and molluscs. White bars represent chromosomes. Vertical lines connect the genomic positions of orthologous genes, colored by the bilaterian ALG to which the genes belong. In both annelids and molluscs, most lineages retain a relatively conserved genome structure characterized by maintenance of ALGs with several lineage-specific chromosome fusion events. Within annelids, the genomes of clitellates (leeches and earthworms) are completely scrambled. Within molluscs, the genome of *Octopus bimaculoides* is highly scrambled.

### Supplemental experimental procedures

#### Assembly selection and acquisition

We first manually curated a dataset of publicly available chromosome-level bilaterian genomes to maximize the number of phyla and classes represented and minimize biases caused by over-sampled taxa. The final dataset consisted of 64 assemblies from 15 bilaterian phyla and at least 52 classes (https://doi.org/10.5281/zenodo.13148652). Assemblies for *Hypsibius dujardini* (phylum Tardigrada) and *Cristatella mucedo* (phylum Bryozoa) were obtained from NCBI Gene Expression Omnibus (GSE169088)[6]. The assembly for *Emplectonema gracile* (phylum Nemertea) was obtained from NCBI BioProject (PRJNA1077883)[7]. Assemblies for *Schizocardium californicum* and *Ptychodera flava* (both phylum Hemichordata) were obtained from Figshare (https://figshare.com/projects/Hemichordate_Genomes/168110)[8]. All other assemblies were obtained from the NCBI Genome database using NCBI Datasets. Many of the assemblies were produced by the Darwin Tree of Life Project[9].

#### Gene prediction

For 38 assemblies, we used RefSeq or GenBank gene models available on the NCBI database. For 15 assemblies, we used gene predictions from published papers that were not available on NCBI[1,4,5,8,10–13]. Gene prediction for the remaining 14 species was performed using a previously published pipeline[4]. Briefly, repeats were first identified with RepeatModeler2 (v2.0.4)[14] and masked using RepeatMasker (v4.1.5)[15], then the BRAKER3 pipeline (v3.0.3)[16] was used for gene prediction. Where available, RNA sequencing data from the same species was used as hints. Protein hints were used if no RNA sequencing data was available. RNA sequencing reads were trimmed with fastp (v0.23.4)[17] and mapped using STAR (v2.7.10b)[18]. BUSCO metazoa *odb10* (v5.4.7)[19] was used to assess the completeness of gene predictions.

#### Phylogenetic analysis and ancestral state reconstruction

OrthoFinder (v2.5.4)[20] was used for orthology assignment. OrthoSNAP (v0.0.1)[21] was used to recover additional single-copy orthologues from gene family trees. MAFFT (v7.520)[22,23], ClipKIT (v1.4.1)[24], and PhyKIT (v1.11.7)[25] were used for sequence alignment, trimming, and concatenation, respectively. The maximum likelihood algorithm of IQ-TREE (v2.1.4)[26] using UFBoot2[27] (1,000 replicates) was utilized for phylogeny construction on a partitioned alignment of 148 concatenated protein sequences with species occupancy of at least 75%. The Markov-chain-Monte-Carlo-based stochastic character mapping approach of Huelsenbeck et al.[28] was used for ancestral state reconstruction. Ancestral state reconstruction was performed using high versus low rearrangement as a discrete character with a unidirectional character state transition model using Ape (v5.7-1)[29] and Phytools (v2.1-1)[30] in R[31].

**Synteny analysis and rearrangement index calculation**

Synteny analysis was implemented using SyntenyFinder[5]. OrthoFinder (v2.5.4)[20] was used to identify genes belonging to bilaterian ALGs based on orthology. Idiogram plots were produced using RIdeogram (v0.2.2)[32]. The rearrangement index (RI) was calculated as described previously[1], where higher indices indicate higher levels of rearrangement. Briefly, the rearrangement index for a given ALG ($R_{ALG}$) is defined as:

$$R_{ALG} = 1 - (S_{CHR} \times C_{CHR}) \quad (1)$$

where $S_{CHR}$ is the ALG splitting parameter (the highest proportion of genes from the ALG on one single chromosome). $C_{CHR}$ is the ALG combining parameter (the proportion of genes on this chromosome that belong to that ALG).

The genome rearrangement index (RI) is then the mean of the rearrangement indices for each ALG in the genome:

$$RI = \frac{\Sigma\,(R_{ALG})}{N} \quad (2)$$

where $N$ is the total number of ALGs. For RI calculations in cnidarian genomes, only ALGs present in the ancestor of cnidarians were considered[3].

**Regression analysis**

We analyzed the relationship between the rate of protein sequence evolution and RI using a logistic regression model. The logistic model was fitted to the data using the non-linear least squares function (nls) in R with the self-starting logistic function (SSlogis).

**Data availability**

The dataset for this study and gene models for the annotated assemblies are deposited on Zenodo (https://doi.org/10.5281/zenodo.13148652). Custom R scripts are available in our GitHub repository (https://github.com/symgenoevolab/animal_genome_structure).

### Author contributions

T.D.L. and Y.-J.L. designed research; T.D.L. performed research; T.D.L., I.J.-Y.L., and Y.-J.L. contributed new reagents/analytic tools; T.D.L. and Y.-J.L. analyzed data; T.D.L. and Y.-J.L. wrote the paper.
